# Targeted Polymersomes Enable Enhanced Delivery to Peripheral Nerves Post-Injury

**DOI:** 10.1101/2024.09.05.611478

**Authors:** Kayleigh Trumbull, Sophia Fetten, Dru Montgomery, Vanessa Marahrens, Olivia Myers, Noah Arnold, Jeffery Twiss, Jessica Larsen

## Abstract

The gold standard therapy for peripheral nerve injuries involves surgical repair, which is invasive and leads to major variations in therapeutic outcomes. Because of this, smaller injuries often go untreated. However, alternative, non-invasive routes of administration are currently unviable due to the presence of the blood-nerve barrier (BNB), which prevents passage of small molecules from the blood into the endoneurium and the nerve. This paper demonstrates that ligands on the surface of nanoparticles, called polymersomes, can enable delivery to the nerve through non-invasive intramuscular injections. Polymersomes made from polyethylene glycol (PEG)-b-polylactic acid (PLA) were conjugated with either apolipoprotein E (ApoE) or rabies virus glycoprotein-based peptide, RVG29 (RVG) and loaded with near infrared dye, AlexaFluor647. ApoE was used to target receptors upregulated in post-injury inflammation, while RVG targets neural specific receptors. Untagged, ApoE-tagged, and RVG-tagged polymersomes were injected at 100 mM either intranerve (IN) or intramuscular (IM) into Sprague Dawley rats post sciatic nerve injury. The addition of the ApoE and RVG tags enabled increased AlexaFluor647 fluorescence in the injury site at 1 hour post IN injection compared to the untagged polymersome control. However, only the RVG-tagged polymersomes increased AlexaFluor647 fluorescence after intramuscular injection. Ex vivo analysis of sciatic nerves demonstrated that ApoE-tagged polymersomes enabled the greatest retention of AlexaFluor647 regardless of the injection route. This led us to conclude that using ApoE to target inflammation enabled the greatest retention of polymersome-delivered payloads while RVG to target neural cells more specifically enabled the penetration of polymersome-delivered payloads. Observations were confirmed by calculating area under the curve pharmacokinetic parameters and the use of a two-compartment pharmacokinetic model.

## Introduction

Severe injury to the peripheral nerves often results in irreversible damage that leaves patients permanently disabled. After the peripheral nerve injury, such as traumatic crushing or severing of the nerve, the axon segments distal to the injury undergo Wallerian degeneration, which is a neuron intrinsic mechanism that results in death of the severed axon with degeneration of their surrounding myelin sheaths to generate microenvironment conducive to axon regeneration^1^. Axon segments proximal to the injury site retract but then begin to regrow in the peripheral nervous system (PNS); however, in the central nervous system (CNS) this spontaneous regrowth rarely occurs^2^. The PNS neurons respond to axonal injury by changing gene expression programs through an injury response supported in part by neurotrophic factors and cytokines that are released by supporting glial cells and inflammatory cells in the nerve. These resulting changes in protein expression support transition to a growth phase to allow for reinnervation of target tissues in the PNS, but these gene expression programs are not seen after CNS injuries. Even though nerves spontaneously regenerate in the PNS, the growth rates are abysmally slow. Consequently, full recovery is successful with injuries with growth distances a few centimeters from the target, but long-range regeneration often needed after nerve injury is rarely successful resulting in permanent disabilities.

The degenerated nerve segment distal to the injury provides a conduit for axons to regenerate, so if the nerve is not completely severed, axons can grow directly into this permissive conduit. Gaps often occur between the proximal and distal nerve stump that make regeneration much less likely without intervention. Consequently, the current gold standard to repair peripheral nerve injuries involves surgical interventions, specifically microsurgery nerve grafts, which are recommended in more severe cases^3,4^. Therapeutic outcomes are greater for patients with injuries over shorter distances, with outcomes declining as distance between distal and proximal stumps increases^5,6^. As an example, examining 26 cases of sports-related axillary (shoulder) nerve injury, patients undergoing surgery with nerve grafts (35%) did experience functional recovery. However, the remaining 65% of patients who did not undergo surgical treatment experienced no recovery^7^. Nerve conduits have also been attempted as therapeutic interventions, being surgically implanted to aid in guiding the regenerating axon toward the distal stump, although they are only recommended for small gaps less than or equal to 3 cm^6^. Increasingly, nerve conduits are being designed with natural or synthetic biomaterial structures to supply the healing nerve with additional therapeutics to improve the healing process. These therapeutics often include growth factors, anti-inflammatory agents, immunosuppressant agents, and voltage-gated potassium channel blockers^8^. Despite their promise, these micro and nanostructures still require an invasive procedure to maximize access to the injured nerves. Clearly, with the wide variation in extent of PNS injuries, surgery is not consistently recommended or consistently successful. However, alternative, non-invasive routes of administration are currently unviable due to the presence of the blood-nerve barrier (BNB). Furthermore, injuries requiring axon regeneration over more than 5-6 cm from to reach target tissues has lower success rates even with surgical intervention because of the slow rates of axon regrowth. In instances requiring such long-distance regeneration, the growth-promoting microenvironment of the degenerating nerve diminishes, and target tissues are less receptive for innervation. Thus, both anatomical repair and growth accelerating interventions are needed.

The BNB is the endoneurial microenvironment that interfaces the neurons and the vasculature of the PNS^9^. It contains tight junctions that serve to protect peripheral nerves by limiting access of harmful or foreign substances, including pathogens, to enter from the bloodstream^10^. This restriction also applies to therapeutics, making treatment of PNS diseases and injuries without surgery incredibly difficult. As such, surgical interventions, either with or without direct injection of a potential therapeutic into the nerve, remain the only clinically viable option for the treatment of peripheral nerve injuries. To circumvent the BNB and enable non-invasive therapy for patients, we have developed targeted nanoparticles, called polymersomes, to deliver payloads to nerves following non-invasive routes of administration.

Polymersomes are vesicles made from amphiphilic diblock co-polymers; for this application, polymersomes are made specifically from polyethylene glycol (PEG, 1000 Da)-b-polylactic acid (PLA, 5000 Da) (PEG-PLA). A hydrophilic ratio roughly between 25-40% leads to the formation of a vesicle^11^ containing a hydrophobic core, capable of encapsulating hydrophobic molecules, an aqueous center, capable of encapsulating hydrophilic drug molecules, and an exterior layer of PEG which helps to shield the vesicle by drawing in water molecules and promoting specific protein adsorption^12,13^ as the polymersomes travel throughout the body. Polymersomes have been utilized previously for the encapsulation of many therapeutic molecules, including peptides, siRNA, proteins, and other macromolecules^14^. Polymersomes are tunable due to the ability to change characteristics such as size and surface charge by manipulating the synthesis parameters^15–18^. They can also be customized by attaching a variety of ligands to the exterior PEG layer to enable targeted delivery of the encapsulated drug.

There is a lack of knowledge on general approaches for the use of nanoparticles in non-invasive delivery to the peripheral nervous system^19^, so researchers turn towards the blood-brain barrier (BBB) as a guide. To passively diffuse through the BBB, polymersomes must be formed at very low diameters, be lipid soluble, and be appropriately charged^20^. Although similar characteristics could allow polymersomes to diffuse through the BNB, this would likely require unreasonably high dosages to generate enough of a concentration gradient to facilitate this diffusion. By attaching ligands to the PEG layer on the surface of polymersomes, we can utilize naturally occurring receptor-mediated transcytosis across the BNB. Apolipoprotein E3 (ApoE) has been shown to bind low-density lipoprotein receptors (LDLR) in the brain, aiding in passage through the BBB^21–23^, a comparable membrane of even higher selectivity than the BNB. However, ApoE may have low specificity for uptake in nerves due to the abundance of LDLR in other organs, including the liver, adrenal, and heart. Rabies virus glycoprotein-based peptide, specifically RVG29 (RVG), shows increased specificity to neurons^24^. RVG is reported to bind nicotinic acetyl-choline receptors (nAChR)^25^ and to neural cell adhesion molecule (NCAM)^26^. In contrast to LDLR, nAChRs and NCAMs are expressed up to 3-fold higher in the brain than in other organs^27^.

In this paper, we developed an understanding of the impact of these two targeting ligands, ApoE and RVG, on the delivery of polymersome payloads across the BNB in the aftermath of a PNS injury. Polymersomes were created and characterized to confirm size distributions and appropriate ligand attachment. After being confirmed to be non-toxic in neural cell lines, the pharmacokinetics of targeted and untargeted polymersomes were assessed using a sciatic nerve injury (ScNI) created in Sprague Dawley rats. Polymersomes encapsulated with a near-infrared dye, AlexaFluor 647 (AF647), without tags or with either ApoE or RVG targeting ligands were injected through both intranerve (IN, current gold-standard) and intramuscular (IM, non-invasive comparison) routes of administration. Animals were assessed over a 48-hour period to determine pharmacokinetic parameters, and post-mortem analysis confirmed safety. We, thus, confirmed the pharmacokinetic differences observed when using these targeting ligands in both direct and non-invasive routes of administration, notably confirming that intramuscular injections can lead to nerve penetration mitigated through targeted polymersomes.

## Results and Discussion

### Polymersome Characteristics

PEG-PLA polymersomes were successfully formed via solvent injection. In general, all particles were of sufficiently small size for injection via a Hamilton syringe. They encapsulated roughly the same amount of AF647, around 2.5% of the total mass loaded, which allowed us to dose for *in vivo* studies using polymer concentration. Untagged polymersomes were consistently formed with an average diameter of 136 ± 5.9 nm, which aligns with previous works in our lab using the same polymer^28^. The consistency in the synthesis of these particles is depicted by the narrow peaks in the intensity-weighted size distributions (Figure 1A) and the relatively low standard deviation. A PDI of 0.04 ± 0.02 confirms that polymersomes are forming with uniform diameters. Figure 1E confirms the vesicular structure of the polymersomes with an expected amount of variability in size for a self-assembled system. The untagged particles have a very negative surface charge of -26.3 mV ± 4.5. Even with their small size, polymersomes cannot passively diffuse through the BNB; their efficiency as a drug delivery vehicle is greatly enhanced by the addition of targeting ligands that utilize receptor-mediated transcytosis to infiltrate nerves^29^. This has the capacity to surpass the efficacy of nerve injury treatment reliant on passive diffusion alone.

**Figure 1.**
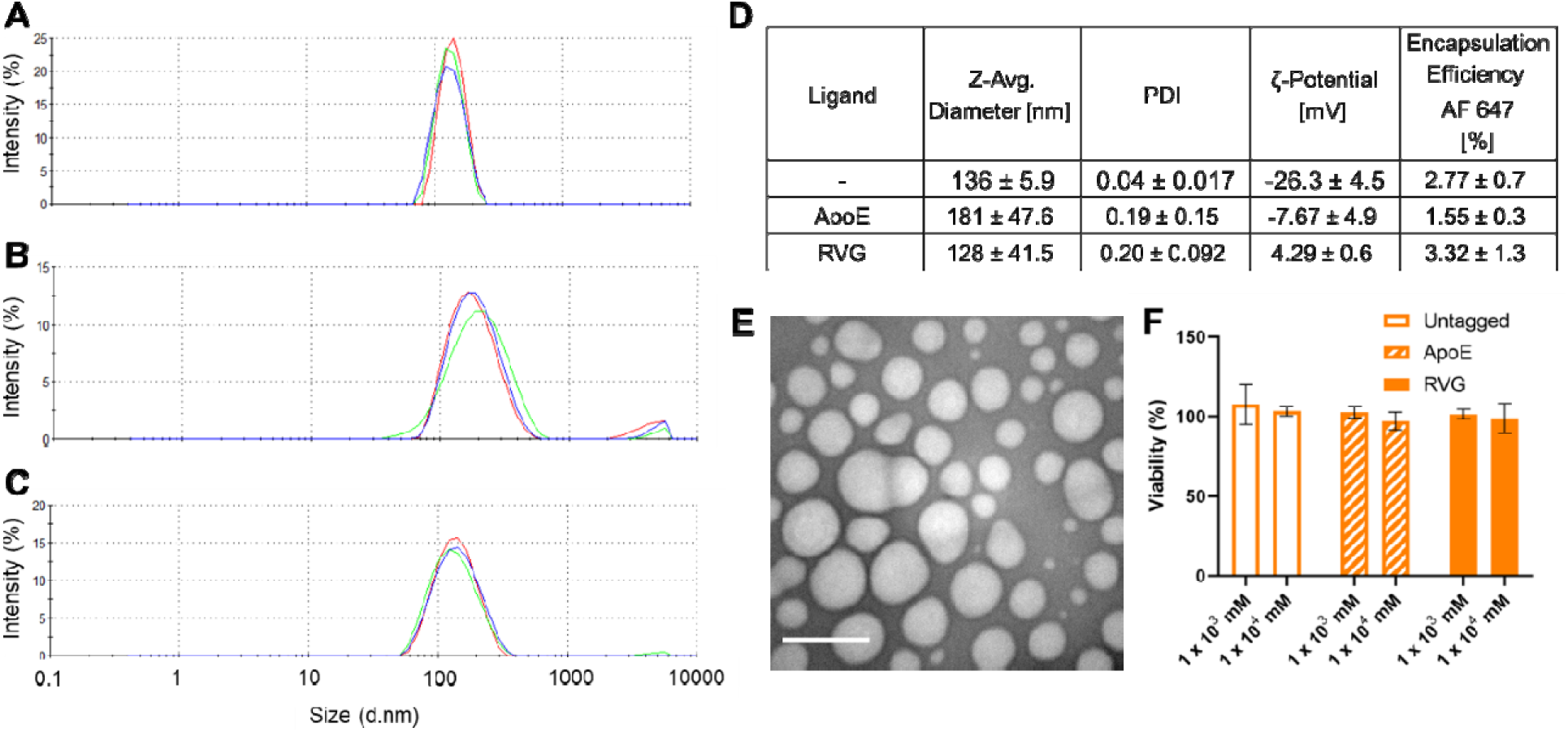
PEG-PLA Polymersome Characterization Data. A. Intensity-weighted size distributions of untagged PEG-PLA polymersomes obtained via dynamic light scattering demonstrate the monodisperse nature of forming polymersomes via solvent injection. Different color tracings represent different runs, highlighting the repeatability of results. B. Intensity-weighted size distributions of ApoE-tagged PEG-PLA polymersomes show the broadening of the overall size distribution and shift to larger sizes caused by the addition of a full-length protein. C. Intensity-weighted size distributions of RVG-tagged PEG-PLA polymersomes show a broadening of the size distribution peak, likely due to the addition of the positive charge. D. The table here highlights the overall properties of polymersomes with various targeting ligands employed in our in vivo studies. E. Transmission electron microscopy (TEM) images of polymersomes demonstrated that most particles had the same diameter, with some overall polydispersity as expected. Scale bar = 200 nm. F. Viability (%) was determined post incubation of PEG-PLA polymersomes in SH-SY5Y cells via an MTS assay at 10 times (10x) and 100x (100x) our in vivo doses. Absorbance measurements determined were compared to untreated cells. No statistically significant decreases in viability were observed.

To assess the successful attachment of the targeting ligands, the same characterization methods are sufficient. ApoE is a larger protein with 299 amino acid residues^30^. Fully uncoiled, this protein would span between 120 and 300 nm. When successfully bonded to the polymersome surface, ApoE will likely maintain a coiled or bent conformation, thus reducing its length^30^. The substantial increase in the diameter of ApoE-tagged polymersomes relative to the untagged polymersome diameter lends to clear identification of successful ApoE binding (Figure 1D). As shown in Figure 1B, the peak of the intensity-weighted size distribution has shifted to the right as the average diameter of the ApoE-tagged polymersomes is 181 ± 47.6 nm. This size shift is accompanied by band broadening and an increase in the PDI of the particles to 0.19 ± 0.15, like what we previously observed^28^. This variability in the particle size is expected due to the size of the ApoE ligand. RVG is a much smaller peptide, spanning around 8 nm, which eliminates the efficacy of comparing particle diameters to verify attachment. However, RVG is a cationic peptide; therefore, successful RVG conjugation can be observed by a shift in polymersome surface charge from negative to positive^31^. RVG-tagged polymersomes have an average surface charge of 4.29 ± 0.6 mV, confirming successful attachment. Although these particles have a similar average diameter to that of the untagged particles, 128 ± 41.5 nm, successful ligand conjugation is also indicated by the widening of the intensity-weighted size distribution in Figure 1C and the PDI of 0.20 ± 0.09 (Figure 1D). The addition of the RVG peptide resulted in an increase in the variability of the polymersomes formed.

When polymersomes with all tags were incubated with SH-SY5Y cells, there was no statistically significant change in viability for treatments of 10 (1 x 10^3^ mM) or 100 (1 x 10^4^ mM) times the dose of polymersomes equivalent to that administered during the *in vivo* studies across the board (Figure 1F). Since these particles are not cytotoxic at such high concentrations, the *in vivo* dosage is also likely to be relatively nontoxic.

### In Vivo Pharmacokinetic Analysis

Following confirmation of *in vitro* safety profiles, Sprague Dawley rats underwent unilateral ScNI followed by either IN or IM injection of untagged (n=3 per route), ApoE-tagged (n=3 per route), or RVG-tagged (n=3 per route) PEG-PLA polymersomes. Sprague Dawley rats were tracked for 48 hours post-ScNI and polymersome injection via an infrared *in vivo* imaging system (IVIS). This enabled the detection of injected, polymersome-encapsulated AF647 in the injury site and throughout the body (Figure 2). In general, IN-injected polymersomes were retained in the injury site at detectible concentrations for at least 4 hours (Figure 2A). Both ApoE and RVG tags led to increased detectible AF647 fluorescence compared to untagged polymersomes. Notably, IM-injected polymersomes resulted in different in vivo behaviors (Figure 2B). The addition of ApoE on the surface of PEG-PLA polymersomes did not appear to increase detectible AF647 signals compared to untagged polymersomes. However, the addition of RVG to polymersomes did lead to detectible AF647 signals remaining for at least one-hour post-injection. Findings were confirmed using region of interest (ROI) analysis of injury sites at each time point (Supplemental Figure 1), which enabled a more robust assessment of the repeatability of findings from animal to animal (Figure 3).

**Figure 2.**
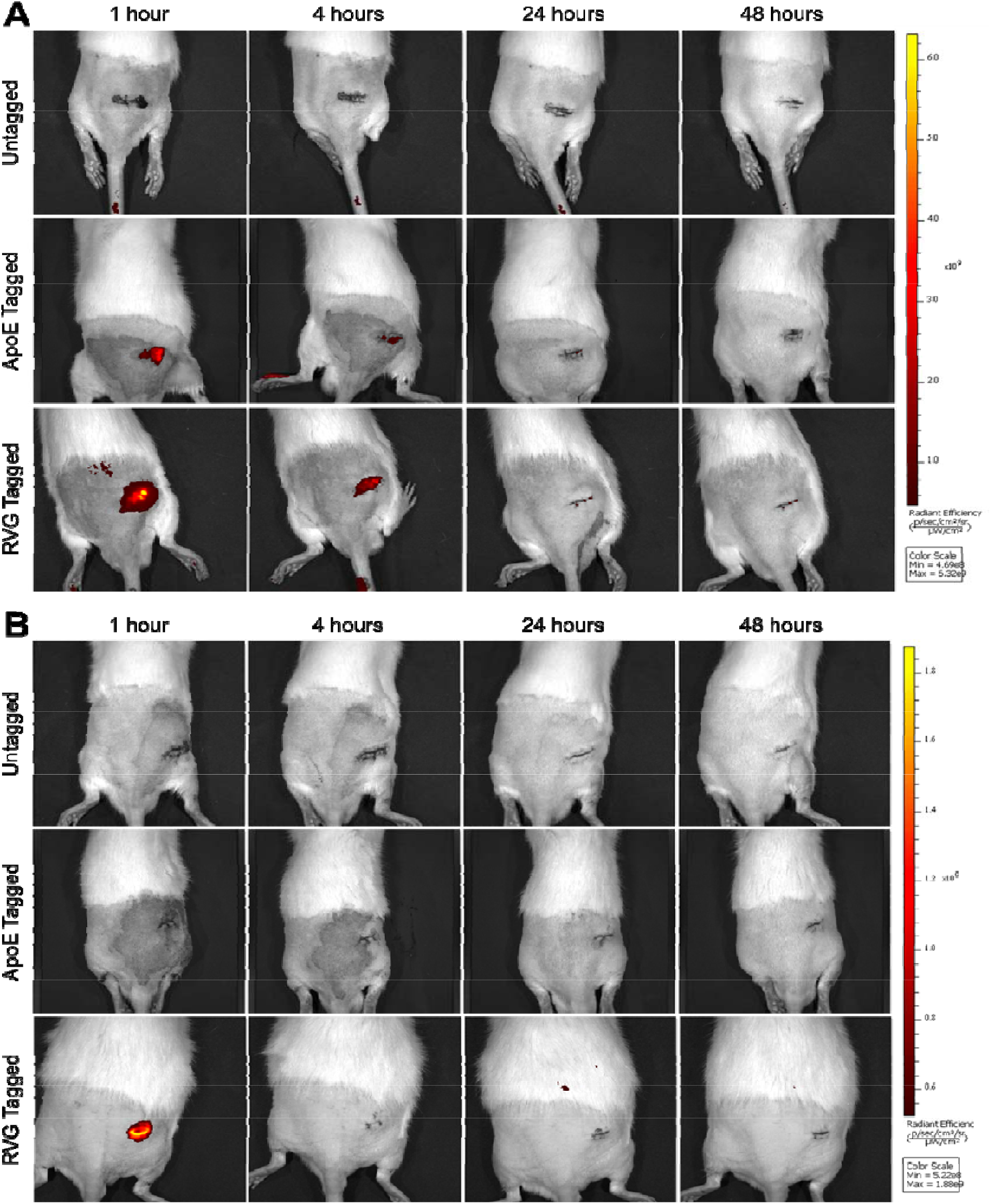
IVIS Imaging of Rats Post Injection. Provided here are example images from one rat per group. A. IN administration led to high retention of PEG-PLA polymersomes over time post injection. Untagged polymersomes were no longer detectible in the injury site only one hour post injection. However, both ApoE and RVG tagged polymersomes were retained for extended periods of time post injection. The scale bar in image A has a range of radiant efficiency from 4.69e8 to 6.32 e9. B. IM administration led to much lower retention of PEG-PLA polymersomes in general compared to IN injected polymersomes. However, the addition of the RVG ligand enabled polymersome detection at one hour post injection. The scale bar in image B has a range of radiant efficiency from 5.22e8 to 1.88e9.

**Figure 3.**
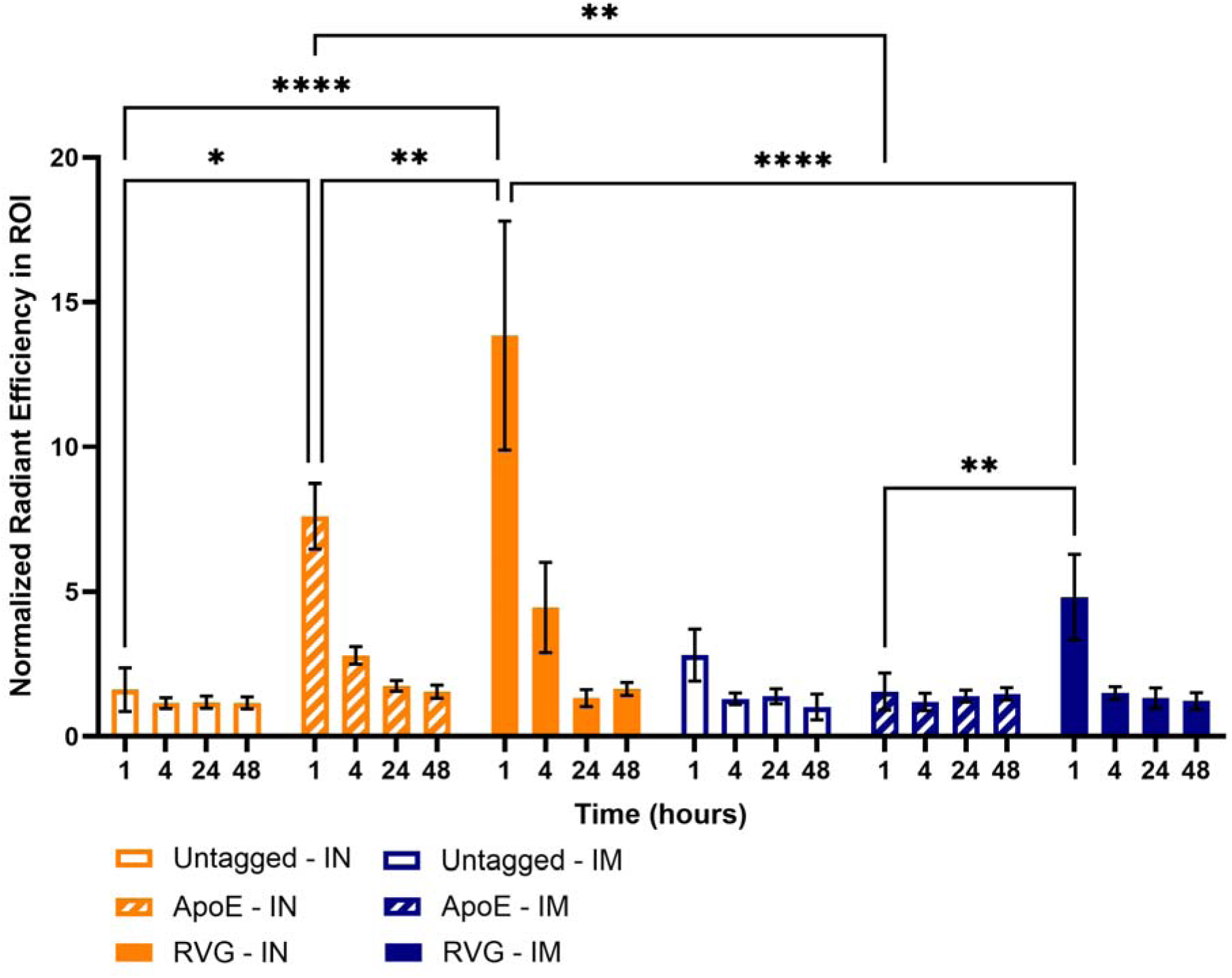
Normalized Radiant Efficiency in the ROI over Time organized by treatment group (n=3 per treatment). In the Living Image software, ROIs were drawn around the injury site to estimate delivery of polymersomes to the site, including a pre-treatment scan for each animal. Measured radiant efficiencies were normalized to pixels of each ROI and then normalized to pre-treatment scans of the same rat. Normalized radiant efficiencies in the ROI between groups at each time point were compared using two-way ANOVA. * p < 0.05; ** p < 0.005, **** p < 0.001

For IN polymersome injections, there were statistical increases in the amount of AF647 fluorescence detected at one-hour post-injection of both ApoE-tagged and RVG-tagged polymersomes compared to untagged polymersomes. This indicates that both tags assisted in the retention of polymersomes near the nerve injection site. It is known that lipoprotein-related proteins like the lipoprotein receptor-related protein-1 (LRP-1) and LDLR are upregulated during inflammation, including in the adult sciatic nerve post injury^32^. Furthermore, this upregulation of LRP-1 has been suggested to play a role in increasing BNB permeability^33^. Therefore, the ability of ApoE, which binds lipoprotein-related receptors, to mitigate increased retention of polymersomes in nerves post-injection is expected. RVG was developed from the neurotropic rabies virus and has been used previously to deliver siRNA across the BBB^34^ due to its ability to bind to the nicotinic acetylcholine receptor^35–37^, which helps us to understand why RVG tagging led to polymersome nerve retention. The largest statistical differences between IN-injected polymersome groups occurred between ApoE and RVG-tagged polymersomes and untagged and RVG-tagged polymersomes. RVG clearly led to the greatest increase in retention of payloads in the nerve for one hour after injury, likely due to its specificity towards neural-specific receptors. It is important to note that despite increased visualization of AF647 via IVIS images with tagged polymersomes at four hours post-IN injection compared to untagged polymersomes (Figure 2A), ROI analysis indicated no statistical differences at any time points other than one-hour post-injection.

As expected, dose-matched IM injections of ApoE and RVG-tagged polymersomes were significantly reduced compared to their IN-injected counterparts. Gold-standard treatment of peripheral nerve injuries involves surgical intervention and/or IN injections. With IM injections, we aim to enable non-invasive treatment routes; as with other extravascular injection routes, we anticipate needing higher doses to reach similar delivery profiles to directly administered particles to account for decreased bioavailability. However, the addition of the RVG-targeting ligand did enable AF647 detection in the injury site at one hour IM post-injection, which was significantly greater than what was observed with ApoE as the targeting ligand or with untargeted polymersomes. As stated above, RVG has specificity to the nicotinic acetylcholine and NCAM receptors, making it more nerve-specific than ApoE. To our knowledge, this is the first time RVG has been used to increase the delivery of nanoparticles to the peripheral nervous system through non-invasive routes of administration.

Pharmacokinetic analysis enables a more in-depth understanding of the absorption, distribution, metabolism, and excretion of polymersomes and enables us to compare across treatment groups. Plasma AF647 concentration versus time curves after injection are presented in Figure 4A-C. As observed in Figures 2 and 3, IN injections led to the retention of polymersomes in or near the injury site. Therefore, over the 48 hours studied, IN injections led to very low concentrations of AF647 in the plasma. This is confirmed by low AUC values for all polymersome types (Figure 4D). IM injections were less retained in the nerve overall and, therefore, led to higher concentrations of AF647 detected in plasma over the time of the study, as observed with all polymersomes regardless of surface tag and confirmed with higher AUC values than corresponding IN injected counterparts. In-depth analysis (Figure 4D) supports IVIS findings and shows that ApoE and RVG lead to very different polymersome delivery behaviors.

**Figure 4.**
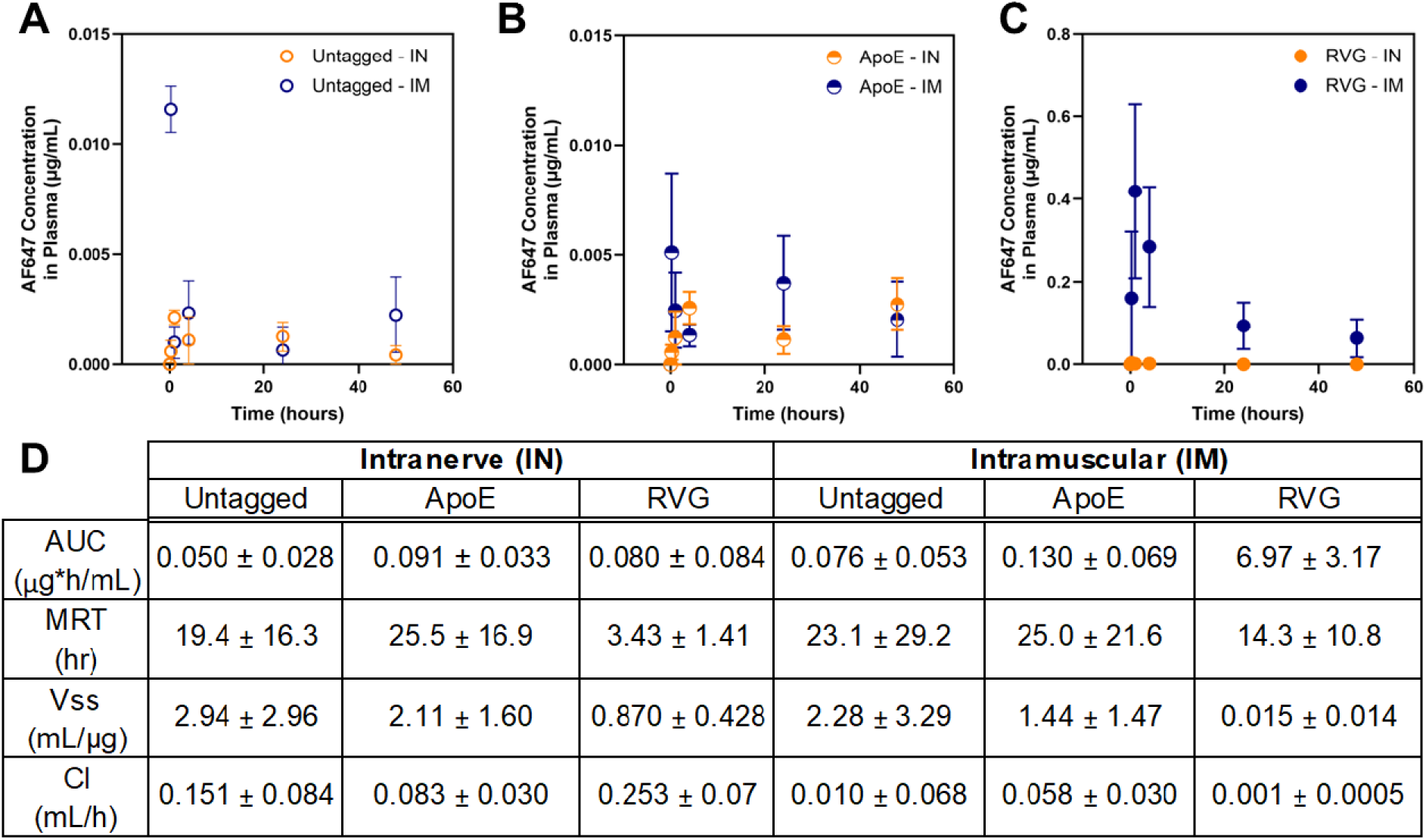
Pharmacokinetic analysis of untagged, ApoE-tagged, and RVG tagged polymersomes injected IN and IM. Blood was collected at various timepoints post injection and plotted on plasma concentration versus time curves. A. Untagged polymersomes vary in plasma concentration of AF647 depending on the injection route, with IM injection leading to rapid plasma presentation of AF647 compared to IN injection, which requires that AF647 leave the nerve before entering the plasma. B. ApoE-tagged polymersomes led to the greatest overall deviation in plasma concentration over time, regardless of IN or IM injection. C. RVG-tagged polymersomes had the highest overall plasma concentration over the duration of the study compared to all other groups. Again, like untagged polymersomes, there was a rapid increase in AF647 plasma concentration with IM injection, and almost no detectible AF647 in plasma after IN injection for the duration of the study. D. This table summarizes the pharmacokinetic data that was calculated based on area under the curve (AUC) of graphs A-C. MRT = mean residence time, V_ss_ = volume at steady state, Cl = clearance.

With regards to IN injections, ApoE outperformed RVG as a targeting ligand to enhance delivery to the nerve. Comparing IN-injected animals, ApoE-tagged polymersomes led to the lowest rate of clearance (Cl) of all tested formulations, with an intermediate volume at steady state (Vss), suggesting an increased residence time within the nerve when analyzed in tandem with IVIS data (Figure 2A and B). ApoE-tagged polymersomes did exhibit a higher mean residence time (MRT) than untagged and RVG tagged polymersomes; however, this is likely an inflated value (with a high deviation) due to the fact that AF647 was clearly not completely excreted during the 48 hour study time (Figure 4B, orange). Further pharmacokinetic studies over longer periods of time will help provide more insights into the MRT for IN-injected polymersomes. IM injections of polymersomes led to increased animal-to-animal variability, demonstrated by higher standard deviations in Figure 4D. However, there are some clear differences in the performance of ApoE-tagged versus RVG-tagged polymersomes when injected IM versus IN, with RVG-tagged polymersomes now appearing to outperform their comparative groups with regard to nerve retention. It is clear that RVG-tagged polymersomes experienced the lowest rate of Cl from the body in the IM injected cohort over the 48 hour time course of the study. This, along with a low Vss and ROI analysis (Figure 3), encourages the conclusion that RVG-tagged polymersomes enable non-invasive delivery to the nerve through IM injections post-ScNI. In general, the MRT of RVG-tagged polymersomes is less than that of untagged polymersomes, whether injected IN or IM, demonstrating that they spend less time in the bloodstream over our study period. Decreased Vss values also indicate that at steady state there are less RVG tagged polymersomes present in plasma compared to their untagged counterparts, meaning that they are spending more time in tissues than both ApoE and untagged polymersomes given that they were injected at the same dose.

Further analysis comparing polymersomes *in vivo* pharmacokinetics using a two-compartment model also highlights major differences (Table 1). Wide confidence intervals were found when modeling IN data, likely due to the low concentrations of AF647 detected in plasma, making it difficult to draw conclusions regarding the significance of findings. However, there are some encouraging potential differences when observed in the context of the body of work. IN-injected ApoE-tagged polymersomes appear to be eliminated at a much slower rate than their untagged and RVG-tagged polymersomes, shown by decreased elimination rate constant (k) values. The ApoE-tagged polymersomes also had the longest calculated half-life (t_1/2_) in the cohort. The data from this model supports what was observed via IVIS (Figure 2A and Figure 3) as well as the AUC-based pharamcokinetic analysis (Figure 4D). With regards to the IM-injected cohort, ApoE-tagged polymersomes had a very large animal-to-animal variability in the AF647 plasma concentration in blood that prevented the establishment of key modeling parameters. We hypothesize that this is due to the nature of ApoE as a targeting ligand. ApoE targets the low-density lipoprotein receptors (LDLR), which are upregulated during inflammatory situations. Therefore, the viability of ApoE as a targeting ligand may vary based on the extent of injury, which can change the inflammatory cascade. Alternatively, RVG targets both NCAM and nAChRs, which are neurally specific; their expression does not depend on the extent of nerve injury^38,39^. However, the RVG-tag on IM-injected polymersomes led to a decreased k_a_ value compared to IM-injected untagged polymersomes, indicating that the RVG tag led to slower absorption of polymersomes into the bloodstream and an elevated k value, indicating that they were removed from the body more slowly. Simultaneously, the t_1/2_ is increased compared to untagged polymersomes, indicating RVG-tagged polymersomes persisted in the body for longer periods of time. By combining these findings with data presented in Figures 2B and 3, we can conclude that the decreased k_a_, increased k, and increased t_1/2_ are likely due to increased nerve retention.

**Table 1.** Pharmacokinetic Parameters calculated by Non-linear analysis of a two-compartment model. Values presented were best fit values with parentheses indicating the 95% confidence interval.

|  | Intranerve (IN) |  |  | Intramuscular (IM) |  |
| --- | --- | --- | --- | --- | --- |
|  | Untagged | ApoE | RVG | Untagged | RVG |
| $k_a$<br>(1/hr) | 0.009409<br>(0.001504 to 0.07900) | 0.022647<br>(0.0005288 to 0.01248) | 0.1713<br>(0.03574 to 1.381) | 3.269<br>(1.439 to ???) | 1.079<br>(0.3630 to 2.687) |
| $k$<br>(1/hr) | 4.886<br>(0.9312 to 42.17) | 0.8645<br>(0.1721 to 4.714) | 33.6<br>(8.955 to 183.8) | 96.89<br>(67.41 to 128.2) | 0.3132<br>(0.09897 to 0.8501) |
| $t_{1/2}$<br>(hr) | 0.1398<br>(0.01620 to 0.7335) | 0.79<br>(0.1449 to 3.967) | 0.02033<br>(0.003715 to 0.07627) | 0.007049<br>(0.005326 to 0.01013) | 2.181<br>(0.8035 to 6.901) |
| $R^2$ | 0.2485 | 0.3045 | 0.5418 | 0.754 | 0.346 |

### Ex vivo analysis

After 48 hours post-ScNI and injection, rats were euthanized, and their organs were collected for post-mortem analysis. Excretory organs, kidneys, livers, and spleens, were collected to get a better understanding of excretion profiles of the various targeted polymersomes. IVIS imaging of excretory organs (Figure 5) demonstrates some key differences in the excretion of IN-injected versus IM-injected polymersomes. In general, excretory organs from animals injected with polymersomes IN, regardless of their targeting ligand, have a higher presence of AF647 compared to their IM-injected counterparts at 48 hours post-injection. Looking at Figure 4A-C, this is likely due to the fact that polymersomes take longer to enter the blood when injected IN versus IM. Polymersomes are also clearly excreted through both the kidneys and the livers, which is expected for nanoparticle delivery, with polymers at less than 5000 Da typically being excreted through the kidney^40^.

**Figure 5.**
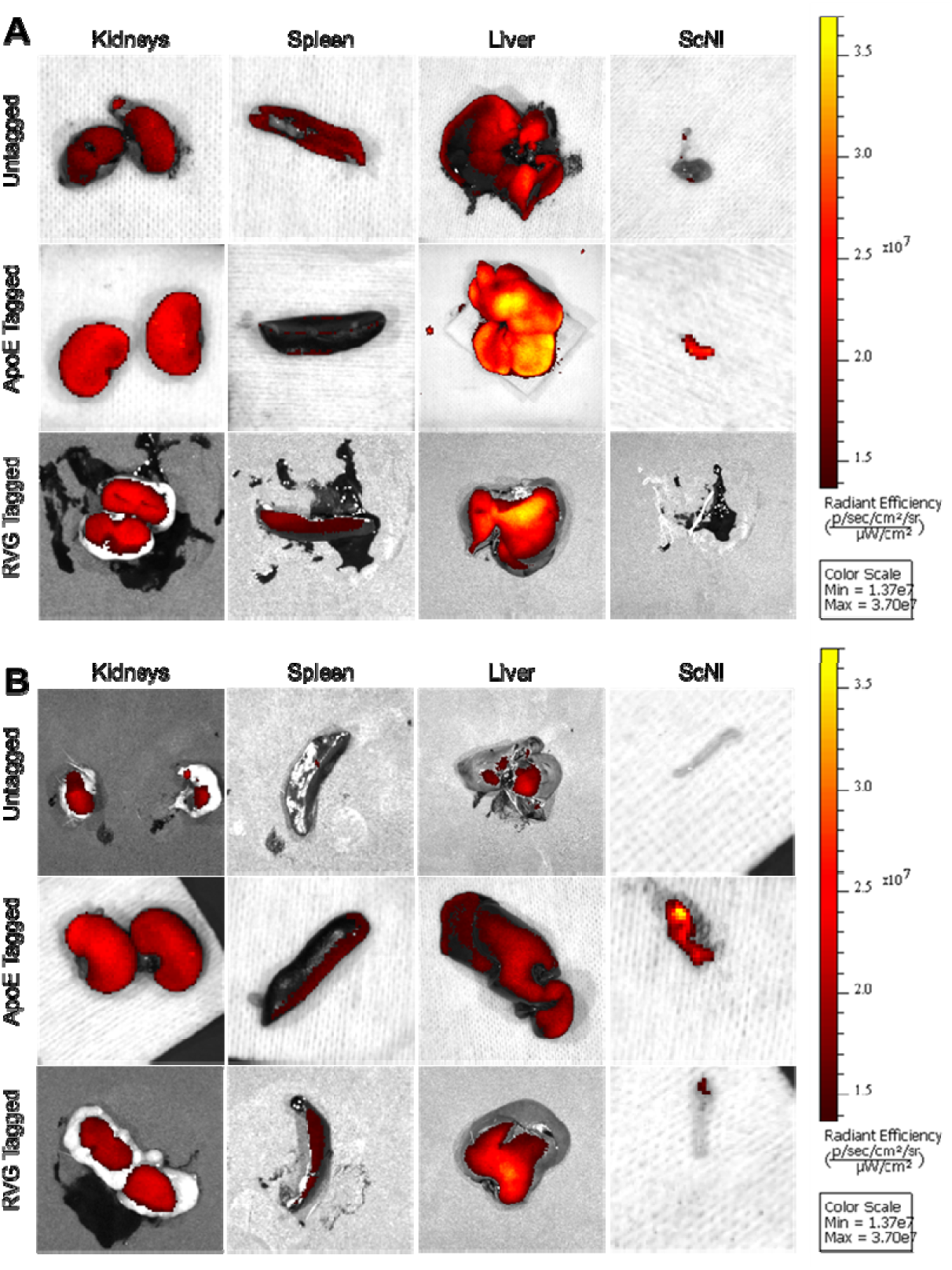
Ex Vivo IVIS images of excretory organs and Sciatic Nerves from one representative animal per treatment group. A. IN injected animals had high AF647 signal in excretory organs post-mortem. B. IM injected animals had lower AF647 signal in excretory organs post-mortem than IN counterparts. The scale bar for images A and B is the same to allow for injection route comparisons. The values for radiant efficiency range from 1.37e7 to 3.70e7.

Additional analysis was performed on sciatic nerves to identify the presence of any remaining AF647 post-mortem using ROI analysis (Figure 6). Statistically, the highest amount of AF647 was observed after injured rats were injected with ApoE-tagged polymersomes via IN administration. This suggests that ApoE leads to a retention of polymersomes in the nerve post-injection for at least 48 hours. This was likely not observed in situ via IVIS due to the limited penetration depth and low sensitivity when imaging very low doses of infrared dye through animal tissue, as the amount near the injury site is clearly decreasing over time based on PK analysis. Ex vivo IVIS images of all rats in this cohort confirm these findings (Supplemental Figure 2A), with RVG-tagged polymersomes and untagged polymersomes injected IN leading to very limited, if any, AF647 expression in sciatic nerves post-mortem. Surprisingly, although not significantly, IM-injected polymersomes may lead to increased AF647 delivery to nerves in 48 hours than their IN-injected counterparts. However, overall post IM injected nerves had a much greater overall deviation in AF647 radiant efficiencies within each treatment group regardless of polymersome tagging (Supplemental Figure 2B). Again, it appears as if ApoE-tagged polymersomes can enable higher retention of injected polymersomes with AF647 based upon the trend towards higher normalized radiant efficiencies in ex vivo nerves, despite whole body IVIS imaging (Figures 2 and 3) suggesting ApoE did not enable an IVIS detectable concentration of AF647 to be delivered after IM injection.

**Figure 6.**
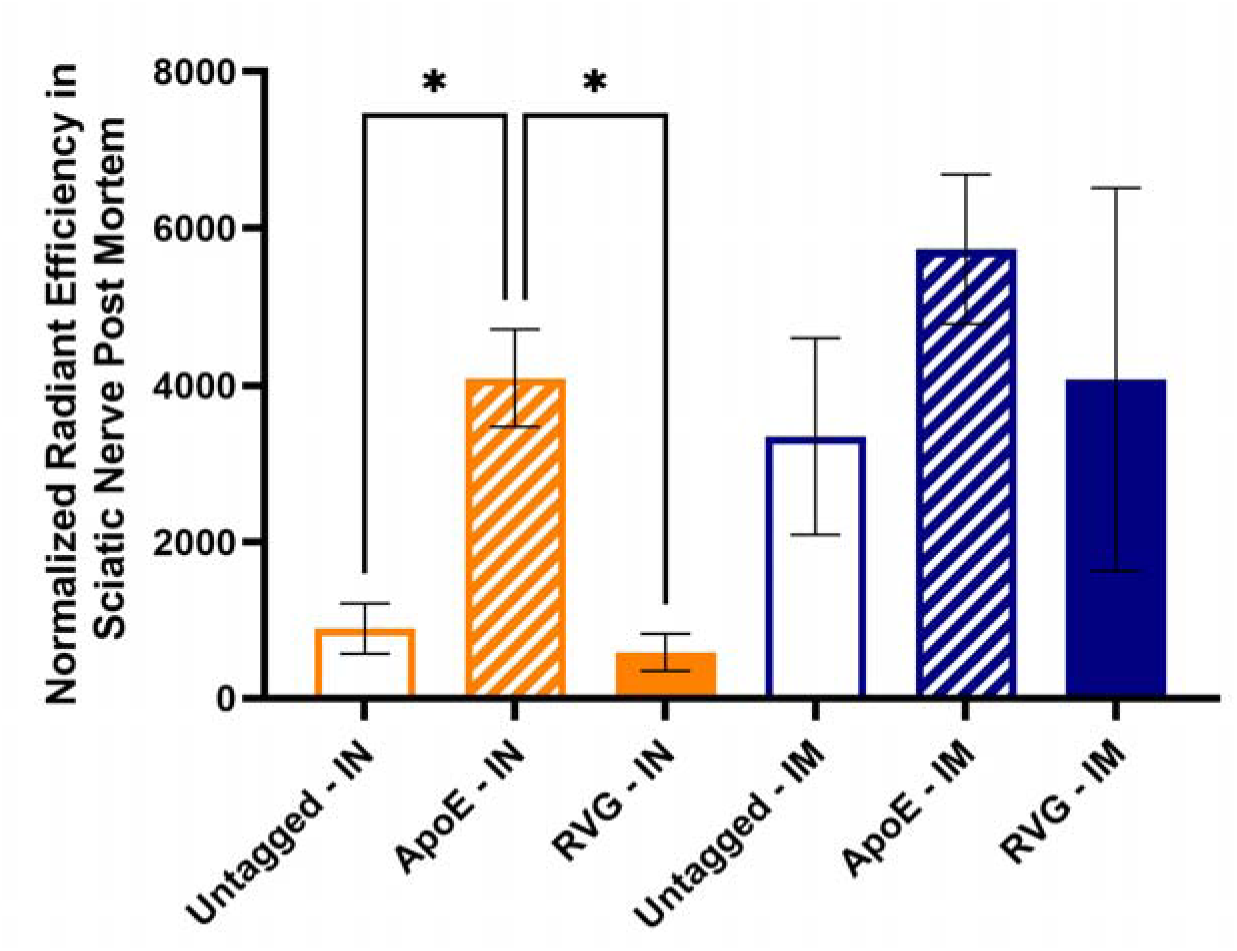
Ex Vivo Region of Interest Analysis on Sciatic Nerves. Results indicate that IN-injected polymersomes with ApoE tags lead to retained AF647 present in the nerve at statistically higher amounts than their untagged and RVG-tagged counterparts. In general, IM injected polymersomes appear to facilitate nerve penetration within the 48-hour time point, as AF647 does appear in postmortem nerves.

Hematoxylin and Eosin (H&E) staining allows for the clear visualization of the microscopic structures of a larger biological tissue and is used to evaluate the health of the excretory organs after processing the injected polymersomes for elimination from the body. The ScN is expected to show tissue damage as a requirement of the *in vivo* nerve injury model, but the tissue excluding the crush site is expected to remain healthy. The ScN images were compared to H&E images of healthy Sprague Dawley rat ScNs showing a resemblance and indicating that the polymersomes themselves did not have any harmful adverse effects on the healthy nerve tissue for each polymersome type and administration route combination^41,42^. Likewise, the spleen, kidneys, and liver were all analyzed *ex vivo* via H&E staining and imaging. Figure 5 shows strong evidence that these tissues all experience substantial contact with the polymersome treatment administered as the body’s innate systems clear and excrete the foreign materials. Figure 7 shows that the excretory organs appeared healthy by the end of the experiment with no harm caused by the presence of the polymersomes regardless of tag ^43–45^. As such, we anticipate that all three polymersomes types are able to be safely and naturally eliminated from the body within a reasonable time period following treatment. These findings were supported by overall maintenance of health of the rats post ScNI and post injection. Rats gained weight at normal rates, continued to groom, and showed no overt signs of toxicity.

**Figure 7.**
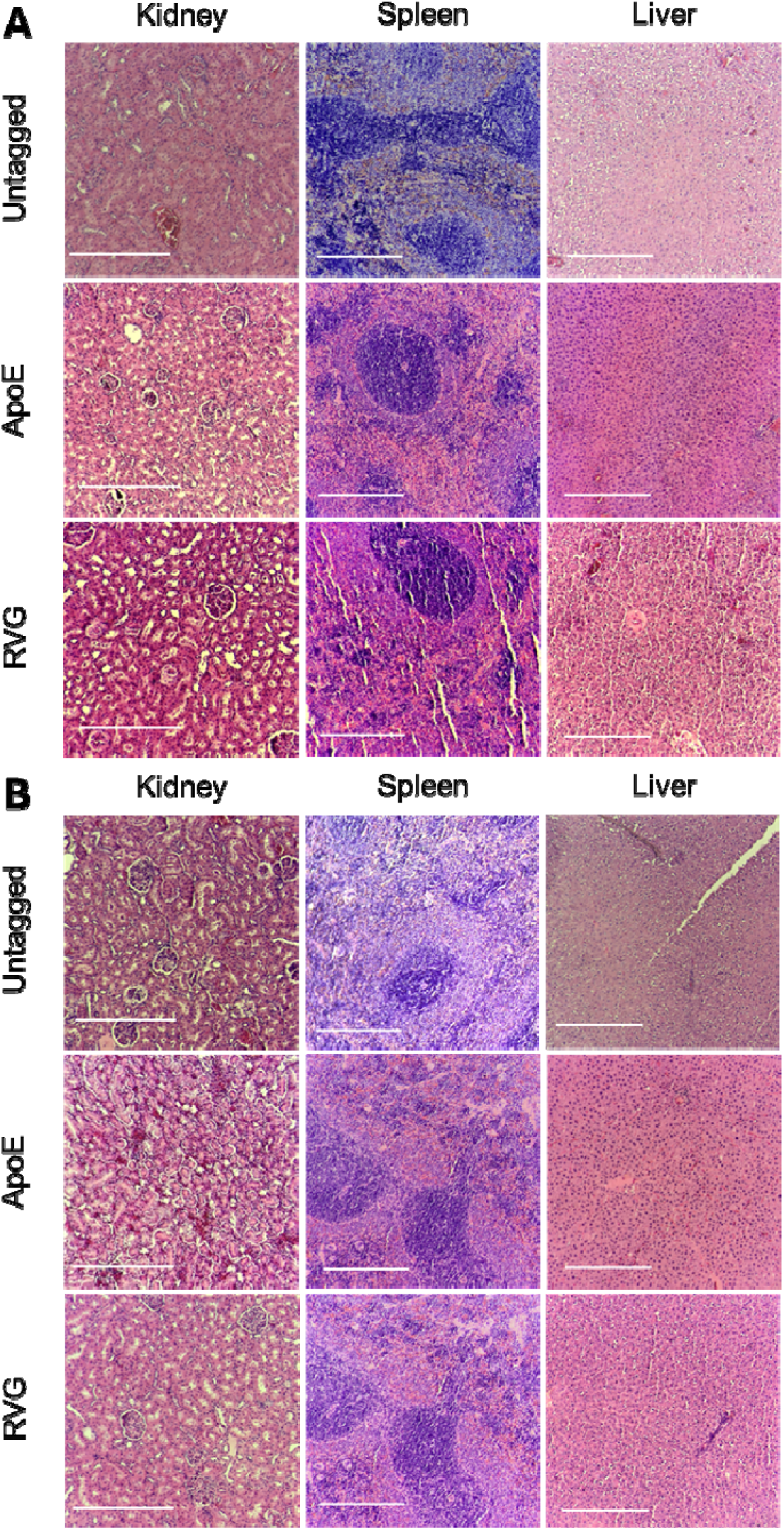
Hematoxylin & Eosin (H&E) staining of excretory organs and Sciatic Nerves from one representative animal per treatment group. A. IN injected polymersomes and B. IM injected polymersomes lead to no observable changes in the health of tissues. Scale bar = 340 µm.

## Conclusions

The information gained from the exploration of the polymersome route of administration and surface targeting ligand combinations presented here offers valuable insight for future considerations of drug delivery to the peripheral nervous system. To maximize future therapeutic effect, polymersomes should be retained in the nerve as long as possible, then cleared through the excretory organs for natural excretion from the body. PEG-PLA polymersomes customized with ApoE or RVG tags were shown to be non-toxic at high doses *in vitro* and cause no undue harm to excretory organs as they are naturally eliminated from the body when injected *in vivo*. Both tags investigated showed increased *in vivo* detection of AF647 1 hour post-IN injection when compared to untagged polymersomes, with ApoE-tagged polymersomes showing more overall retention in the body for the 48-hour study supported by IVIS and pharmacokinetic analysis. Longer *in vivo* studies could improve our understanding of the pharmacokinetic properties of these polymersomes, especially by allowing us to capture the full MRT and half-lives of IN-injected polymersomes. When administered via IM injection, the addition of RVG significantly increased retention of AF647 1-hour post-injection over ApoE-tagged and untagged polymersomes, supported by both IVIS and pharmacokinetic findings. However, it is interesting to note that although AF647 was not detectible after IM injections of ApoE-tagged polymersomes during whole-body in vivo analysis, ex vivo images demonstrated some nerve retention. It appears as if the RVG-tag assists in nerve penetration, but the ApoE-tag may assist in increased nerve retention after injection. Future dose escalation studies may provide additional insight into the efficacy of IM-injected polymersomes and the effect of the RVG tag. Increasing the size of the IM-injected cohort for ApoE-tagged polymersomes may also help to clarify or confirm the animal-to-animal variability in nerve retention. These results are encouraging for the future development of novel nerve injury treatments that are less invasive than current gold-standard treatments as these results show promise for a potential noninvasive, IM injection option for patients suffering from peripheral nerve injuries.

## Experimental Section

### Polymersome Synthesis via Solvent Injection Method

The polymer polyethylene glycol (1000Da)-b-poly lactic acid (5000Da) (Polysciences, Warrington, PA) (PEG-PLA) forms nanoparticular vesicles via self-assembly. A polymer solution of 5mg/mL concentration was formulated by weighing 0.5mg of PEG-PLA into a microcentrifuge tube and dissolving in 100μL of dimethyl sulfoxide (DMSO). Solvent injection method was used with syringe pump speed of 10 μL/min, syringe diameter of 3.58625 mm (½ mL syringe), and 20-gauge needle. The polymer solution was injected into 10 mL of a stirring solution of 2 wt% mannitol in type I deionized (MilliQ) water, which is a standard protocol for the Larsen Lab^46–50^.

### ApoE Conjugation

The previously detailed solvent injection method is consistent with the addition of the reacting groups that bond the ligand. Adding 10mg NHS-PEG(2000)-NHS (JenKem Technology, Plano, TX) to the polymer solution extends the PEG strand. An aliquot of ApoE in 5mM sodium phosphate buffer is added to the mannitol solution. ApoE is attached through the formation of an amide bind (Figure 8A).

**Figure 8.**
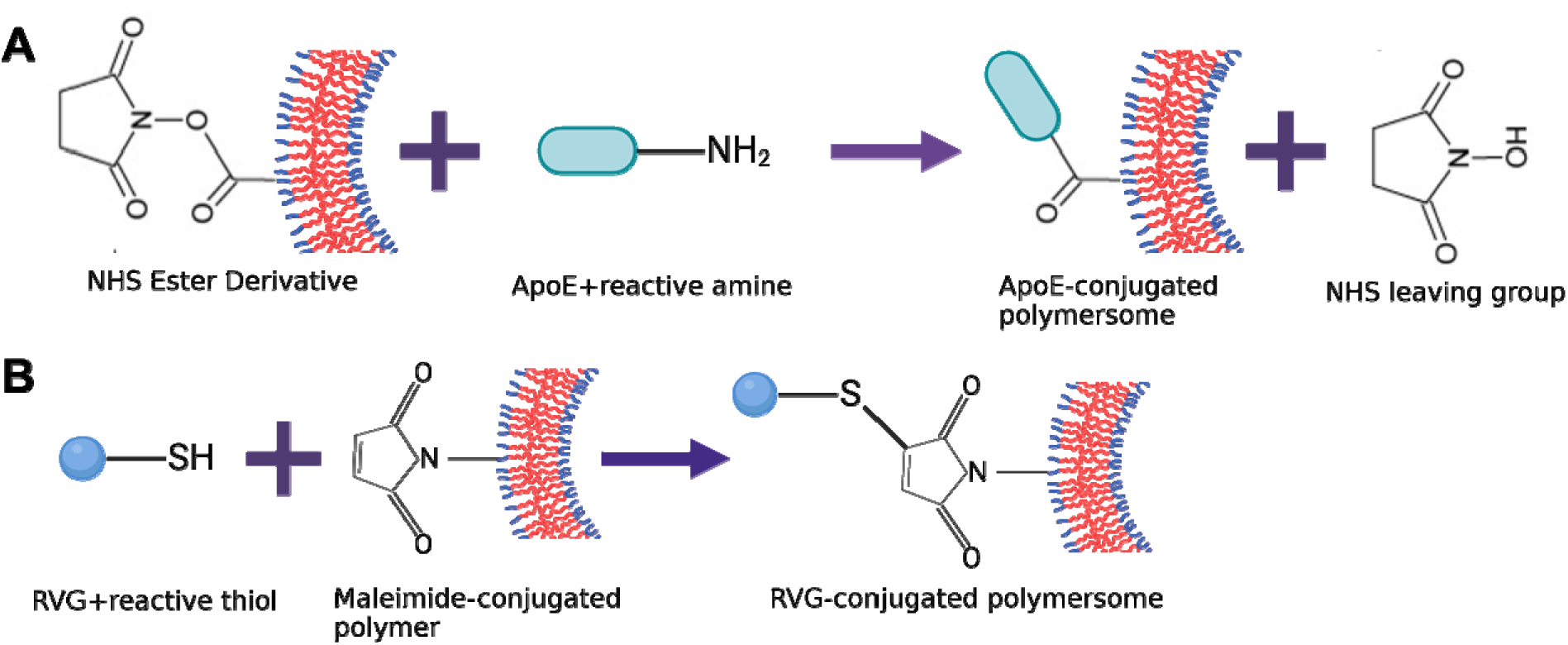
Bioconjugation chemistry used to attach A. ApoE and B. RVG to polymersome surfaces.

**Figure 9A.**
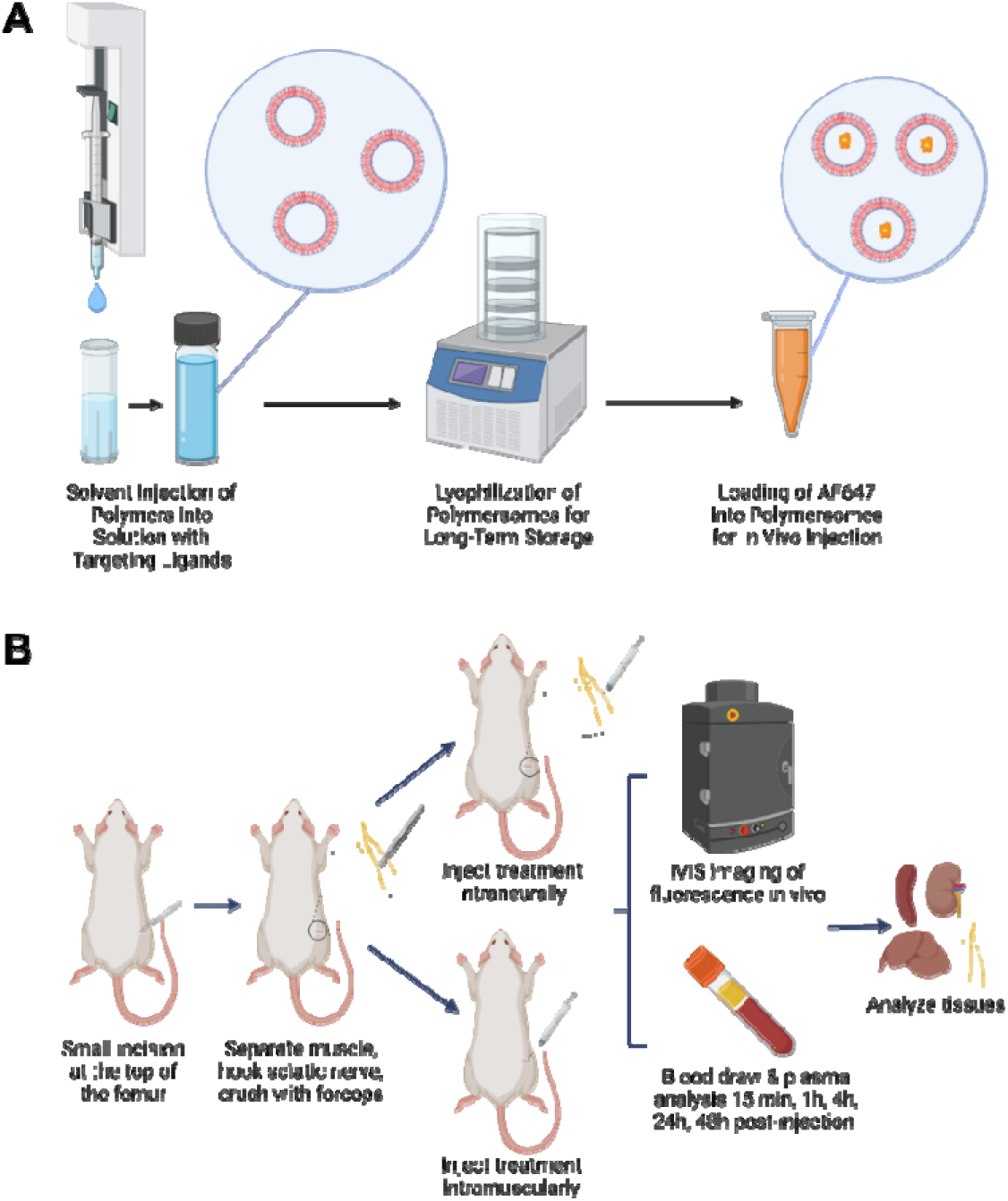
Schematic of polymersome preparation methods, B. Schematic of polymersome injection and in vivo assessment.

### RVG Conjugation

Instead of just PEG-PLA, 0.5mg PEG(1000)-PLA(5000) with a maleimide conjugation (PEG-PLA-MAL) (Broadpharm, San Diego, CA) is measured in a separate microcentrifuge tube and dissolved in 100μL DMSO, then 50μL of each polymer solution is combined in a new microcentrifuge tube. This creates a solution containing an optimized 50:50 mixture of PEG-PLA and PEG-PLA-MAL. Cysteine-conjugated RVG29 is custom ordered from GenScript (Piscataway, NJ) and concentrated at 10mg/mL. With the addition of 50μL of the RVG solution to the mannitol solution, RVG is attached via a reaction between the maleimide conjugated block copolymer and the thiol group in the Cys-conjugated RVG (Figure 8B).

### Lyophilization for Long-Term Storage

Each batch of polymersomes was moved to a 50mL Falcon tube and frozen at -20°C for a minimum of 2 hours with the tube oriented horizontally to minimize crystal formation^15^. The polymersomes were then moved to a -80°C freezer for a minimum of 2 hours before being freeze-dried. Polymersomes were lyophilized in a Labconco FreeZone 4.5L -105C Benchtop Freeze Dyer set to 0.000mbar and -105°C (Kansas City, MO). Lyophilized polymersomes are stored in a desiccator at room temperature.

### Polymersome Loading and Characterization

After synthesis, the average diameter and ζ-potential of the polymersomes was assessed with dynamic light scattering (DLS) using Malvern Instruments’ Zetasizer Nano ZS90 (Malvern, UK). PEG-PLA polymersome morphology was verified using transmission electron microscopy (TEM). TEM images were taken using a Hitachi 7830 UHR transmission electron microscope (Tokyo, Japan). Encapsulation efficiency (EE) of the NIR dye, AlexaFluor 647 (AF647) was determined on a mass basis. A 10mg/mL solution of AF647 Carboxylic acid tris(triethylammonium) salt (Invitrogen, Waltham, MA) was prepared by dissolving 1 mg AF647 in 100μL dimethylformamide. 10mg of lyophilized polymersomes were weighed into a microcentrifuge tube, then 5μL of AF647 solution and 995μL MilliQ water was added to reconstitute the polymersomes by vortexing briefly to mix (Figure 8A). The loaded polymersomes were dispensed into two 100kD microcentrifuge filter tubes (Millipore Sigma, Burlington, MA) and centrifuged for 15 minutes at 14,000xg. Then, the filters were flipped upside down into new microcentrifuge tubes containing 200μL each of DMSO and centrifuged again for 15 minutes at 14,000xg. The dissolved polymersome solution was transferred to a black 96-well plate in duplicates and fluorescence intensity was measured using a BioTek Synergy H1 microplate reader (Aligent Technologies, Santa Clara, CA) and accompanying Gen5 software (Aligent Technologies, Santa Clara, CA) with excitation and emission wavelengths set to 650 and 665nm, respectively.

### Polymersome Preparation for in vitro use

Polymersomes concentrated for *in vitro* use were prepared in an equivalent dose based on the 100mM concentration used for *in vivo* dosage. This concentration amounts to 6mg of polymersomes in each dose administered to a rat weighing 234g, on average. A body weight to total body surface area ratio of 219.2g/356.6cm^2^ for Sprague Dawley rats was used to scale the dose to a 1cm^2^ well on a 96-well plate^51^. This equates to a dose of 0.0158mg/cm^2^. Enough lyophilized polymersomes to make a 10x dose were weighed into a microcentrifuge tube in excess and the appropriate volume of SH-Sy5Y cell media was added so that each 0.0158mg dose is contained in 20μL volume. The dose solutions were pushed through a Whatman UNIFLO 0.45μm syringe filter with 0.45μm pores (Cytiva, Marlborough, MA) to sterilize and stored at -20°C until needed.

### MTS Assay

SH-SY5Y cells (American Type Culture Collection, Manassas, VA) were seeded at a density of 10,000 cells per well in a 96-well plate and incubated overnight prior to treatment. After removing old media and washing cells with 0.1M phosphate buffered saline (PBS), each well was treated with 20μL of the respective polymersome treatment and 180μL of fresh media to replenish the nutrient supply of the filtered media. The cells were incubated for 18-24 hours, then 20μL of MTS Cell Proliferation Assay (Promega, Madison, WI) was added. Between 3-4 hours after adding the MTS reagent, the absorbance of the cells was measured using the microplate reader. The absorbance measurements were normalized to the media only control group to assess relative viability.

### Polymersome Preparation for in vivo use

In a microcentrifuge tube, 10mg of lyophilized polymersomes were weighed, then 5μL of the AF647 solution and 995μL MilliQ water was added to reconstitute the polymersomes by vortexing briefly to mix. The loaded polymersomes were dispensed into two 100kD microcentrifuge filter tubes (Millipore Sigma, Burlington, MA) and centrifuged for 15 minutes at 14,000xg. Then, the filters were flipped upside down into new microcentrifuge tubes and centrifuged again for 15 minutes at 14,000xg. The volume of collected product was determined using micropipettes and the process was repeated until enough polymersomes were collected to make sufficient 100mM polymer-based doses. PBS was added to properly concentrate the solution, ensuring preparations of appropriate dosing volumes for rats of 10 µL.

### Sciatic Nerve Injury (ScNI)

Animal experiments were performed with approval from Clemson University’s Institutional Animal Care and Use Committee (IACUC) through Animal Use Protocol 2023-0105. A total of 18 Sprague Dawley adult rats (mixed M and F) underwent survival surgery with unilateral sciatic nerve forceps crush injuries being performed on their right side in accordance with established protocols^52^. Following ScNI, polymersomes were either administered intraneural (IN), directly into the injury site, or intramuscularly (IM), near the injury site using a 32-gauge needle affixed to a 10μL Hamilton syringe (World Precision Instruments, Sarasota, FL). For treatments administered intranervously, the syringe was inserted into the nerve at a 45° angle and half of the treatment was injected. The needle was repositioned prior to injecting the remainder of the treatment dose. The nerve was released, and sutures applied to close the wound. For treatments administered intramuscularly, the nerve was released after the crush injury was made. The same needle and syringe combination was used to inject the treatment into the muscle near the injury site in two places.

### Live Animal Data Collection

After the ScNI was complete, a blood sample was collected using the lateral saphenous vein at 15 minutes, 1 hour, 4 hours, 24 hours, and 48 hours post injection. The blood samples were frozen at -20°C until they were ready for use. Whole body IVIS images were taken using the IVIS Spectrum Fluorescence and Bioluminescence Scanner (PerkinElmer, Inc., Hopkinton, MA) at 1 hour, 4 hours, 24 hours, and 48 hours post-injection. The study was concluded after 48 hours. The excretory organs (spleen, liver, and kidneys) and sciatic nerve were harvested and imaged via IVIS before being fixed in a 4% paraformaldehyde solution for 24-48 hours at 4°C. The tissues were then moved to a 30% sucrose in PBS solution for long-term storage at 4°C.

### Plasma Analysis

The blood samples were thawed and centrifuged at 9,000xg for 5 minutes. The opaque plasma was transferred in 2μL duplicates into the wells of a BioTek Take3 Microvolume plate (Aligent Technologies, Santa Clara, CA). The fluorescence intensity of the plasma was measured in the microplate reader, for the Take3 plate. The mass of AF647 in the blood was calculated using a calibration curve of known AF647 concentrations in untreated Sprague Dawley rat plasma. Concentration of AF647 versus time plots were created for each polymersome treatment group and injection route. These curves were used to determine pharmacokinetic parameters: Area under the Curve (AUC) and Area under the Moment Curve (AUMC) values were determined using GraphPad Prism 10. From AUC and AUMC, values for Mean Residence Time (MRT), Volume of Distribution at Steady State (V_ss_), and Clearance (Cl) were determined using standard equations for extravascular administration^53,54^. GraphPad Prism 10 was used to perform nonlinear analyses to determine first order absorption rate (ka), first order elimination rate (k), and half-life (t1/2) assuming a two-compartment model, where the nerve is the first compartment, and the rest of the body is the second compartment^55^.

### IVIS Data Analysis

For each rat, a pre-scan was performed to established base line fluorescence of each animal with the excitation and emission wavelengths of AF647. For appropriate comparisons over time, rats were combined in one stack using the Living Image version 4.7.4 software (PerkinElmer, Inc., Hopkinton, MA) to be placed on the same radiant efficiency scale. Regions of interest (ROIs) were established on the right hindlimb near the injury site (Supplemental Figure 1) to track radiant efficiency over time. Measurements from ROIs were normalized to their area in pixels, as the injury site does not show up in the same location for each image, and to the corresponding rat pre-treatment scan to calculate the normalized radiant efficiency values reported.

### H&E Staining of Excretory Organs and Sciatic Nerve Tissues

After fixation and sucrose wash, spleen, liver, kidneys, and sciatic nerve tissues were cut into small pieces (a few centimeters) and placed into tissue embedding cassettes. The samples were dehydrated, and paraffin embedded using the 70% EtOH cycle on a Leica ASP300 S Tissue Processor (Leica Biosystems, Wetzlar, Germany). The samples were then set in paraffin molds using a TEC-II Tissue Embedding Center Dispensing Console and Cold Plate Console (General Data Healthcare, Cincinnati, OH) and allowed to set overnight. The samples were sectioned using a Leica RM 2155 Microtome (Leica Biosystems, Wetzlar, Germany) and set on microscope slides using a water bath at 40°C. The slides were placed on a slide warmer at 40°C for 25-30 minutes to set, then allowed to rest overnight prior to staining. A Hematoxylin & Eosin (H&E) staining (Richard-Allan Scientific, San Diego, CA) protocol suitable for paraffin was used to stain the slides. Mounting media and coverslips were applied, then allowed to dry prior to imaging. The sections were imaged using an Echo Revolve Microscope (Echo, San Diego, CA). The spleen, liver, and kidney sections were imaged at 10x magnification and the sciatic nerve sections were imaged and 10x and 20x magnifications.

### Statistical Analyses

Statistical analyses were performed for each experiment in accordance with most appropriate statistical tests. Normalized radiant efficiency in the region of interest (ROI) measurements and MTS data was analyzed using two-way ANOVA with Tukey test to correct for multiple comparisons with a threshold p-value of 0.05. Radiant efficiencies in post-mortem sciatic nerves were compared within injection groups (IN or IM) using a Brown-Forsythe and Welch one-way ANOVA tests, using Dunnett T3 for correction of multiple comparisons with a threshold p-value of 0.05. All statistical analyses were performed using GraphPad Prism 10.

## Acknowledgements

We would like to thank undergraduate researchers who laid the foundation for this work through their early efforts towards this project: Conner Lumb, Austin Evers, Cheyenne Brady, Jessica Kellner, Valerie Zawrotuk, and Nicholas Johnson. We would also like to thank Lucian Williams for his assistance with slide preparation. Elizabeth Thames and Sharmina Miller provided support and training on the sciatic nerve injury model. Finally, Travis Pruitt and all of the staff at Godley Snell provided assistance with animal husbandry and procedures.

