## Supplemental Figures for "Targeted Polymersomes Enable Enhanced Delivery to Peripheral Nerves Post-Injury"

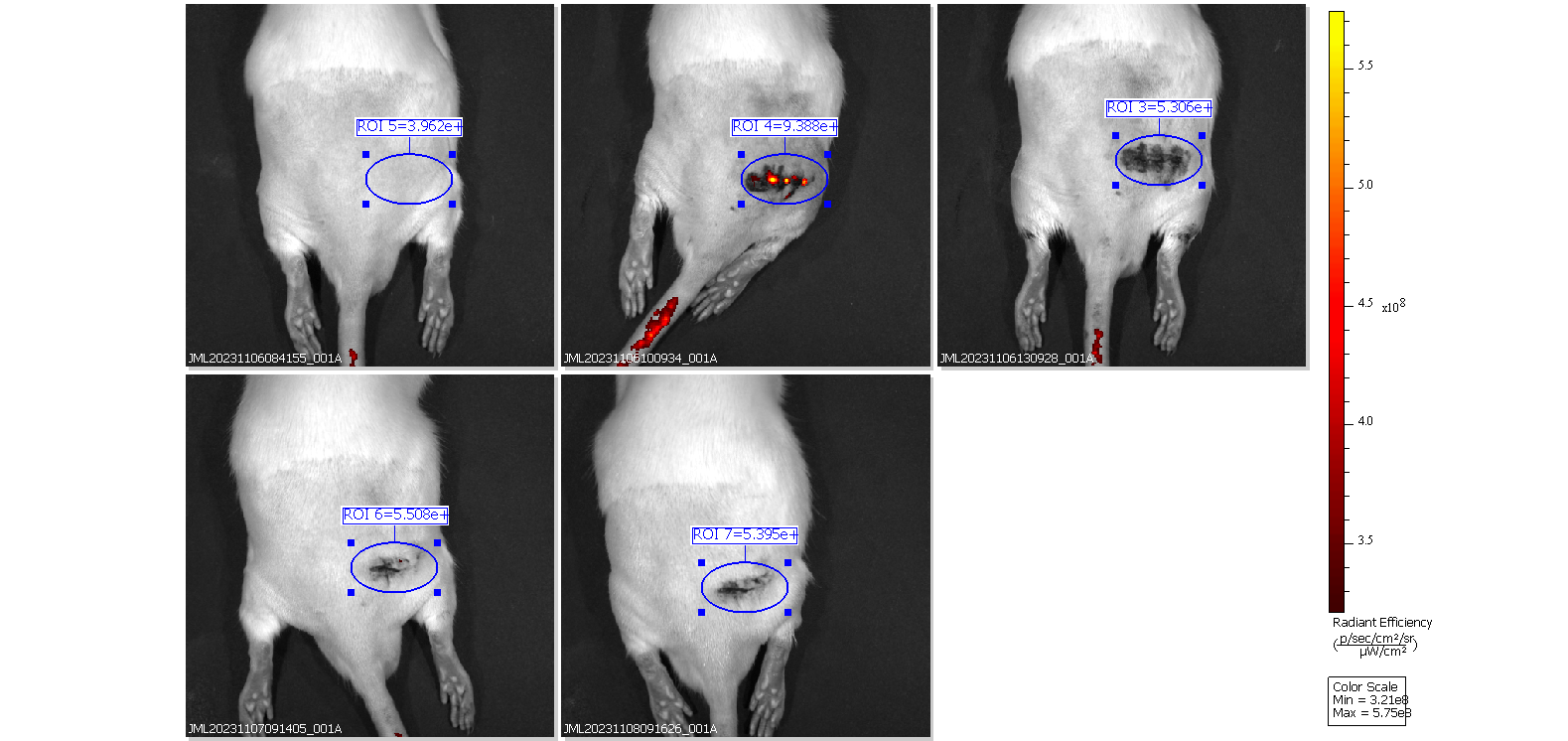

**Supplemental Figure 1. LivingImage Enables ROI Analysis. Here** is an example of region of interest (ROI) analysis of rats using preinjury (left corner) and post-injury analysis of the sciatic nerve injury site. Surgical sites are used to direct analysis location. All ROIs are normalized to each individual rat pre-injury and to changing size of ROI, measured via pixels.

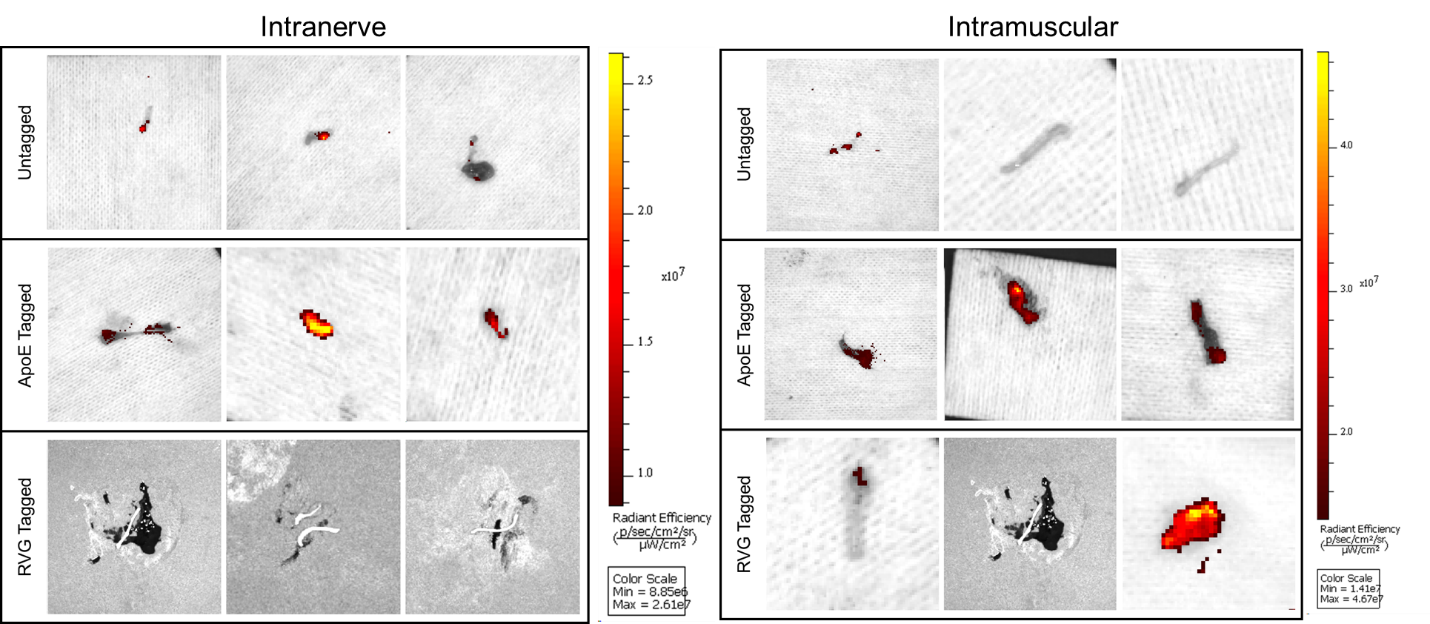

**Supplemental Figure 2. Ex Vivo IVIS images of Sciatic Nerves at 48 hours post-injection.** Sciatic nerve ex vivo analysis demonstrates retention of AF647 in nerves post IN injection with both untagged and ApoE-tagged polymersomes. However, RVG-tagged polymersomes are not retained at detectible fluorescence levels. Post IM injections, ApoE-tagged and RVG-tagged polymersomes are partially retained, with animal-to-animal variability.
